# Mesenchymal Stem Cell Membrane Coating of Gelatin Nanospheres with Drug Release Ability for Inflammatory Tissue Targeting

**DOI:** 10.1101/2025.06.01.657318

**Authors:** Keisuke Moriyama, Mitsuru Ando, Kazuo Sakurai, Isamu Akiba, Yasuhiko Tabata

**Affiliations:** Laboratory of Biomaterials, Institute for Life and Medical Sciences, Kyoto University, 53 Kawara-cho Sohgoin, Sakyo-ku, Kyoto, 606-8507, Japan; Department of Chemistry and Biochemistry, The University of Kitakyushu, 1-1 Hibikino, Kitakyushu, Fukuoka, 808-0135, Japan; Faculty of Environmental Engineering, The University of Kitakyushu, 1-1 Hibikino, Wakamatsu, Kitakyushu, Fukuoka 808-0135, Japan; Department of Plastic and Reconstructive Surgery, Graduate School of Medicine, Kyoto University, 54 Kawara-cho Sohgoin, Sakyo-ku, Kyoto, 606-8507, Japan

**Keywords:** Gelatin nanosphere, Mesenchymal stem cell, plasma membrane coating, controlled release, inflammatory tissue tropism

## Abstract

Improvements in drug delivery have been achieved using nanospheres to prolong drug efficacy, accelerate absorption, and target tissues with ongoing inflammation. Although nanospheres have numerous pharmacokinetic advantages their tissue-targeting ability is poor. In contrast, mesenchymal stem cells (MSC) accumulate in high numbers in inflammatory tissues via the interaction between CXC-chemokine receptor 4 (CXCR4) expressed on MSC and stromal cell-derived factor 1 (SDF-1) secreted during inflammation. Therefore, this study investigated coating gelatin nanospheres (GNS) with MSC membranes (MSC-GNS) by extrusion and ultrasonication methods to enhance their inflammatory tissue tropism. ζ-potential measurements, western blotting and single-particle analysis of MSC-GNS by flow cytometry demonstrated the GNS surface was successfully coated with MSC membranes. Dot blotting demonstrated the binding ability of CXCR4 for SDF-1 was retained by MSC-GNS but absent for MSC-membrane-free GNS. The blood clearance of MSC-GNS was examined by their intravenous injection into mice. Although MSC-GNS and GNS were retained during the early distribution phase, MSC-GNS had a higher retention than GNS during the later elimination phase. Finally, we investigated the tissue distribution of MSC-GNS by intravenous injection into a mouse model of liver fibrosis and their potential therapeutic effect on liver fibrosis. We found a higher accumulation of MSC-GNS in inflamed livers and higher blood retention compared with MSC-membrane-free GNS.

Furthermore, MSC-GNS loaded with an anti-fibrotic agent (LSKL, a 4-amino acid peptide that inhibits fibrosis progression) had an enhanced therapeutic effect on liver fibrosis than uncoated nanoparticles. Therefore, MSC-GNS might be a drug carrier with inflammatory tissue targeting and controlled drug release abilities.

**Synopsis Table of Contents Graphic:** 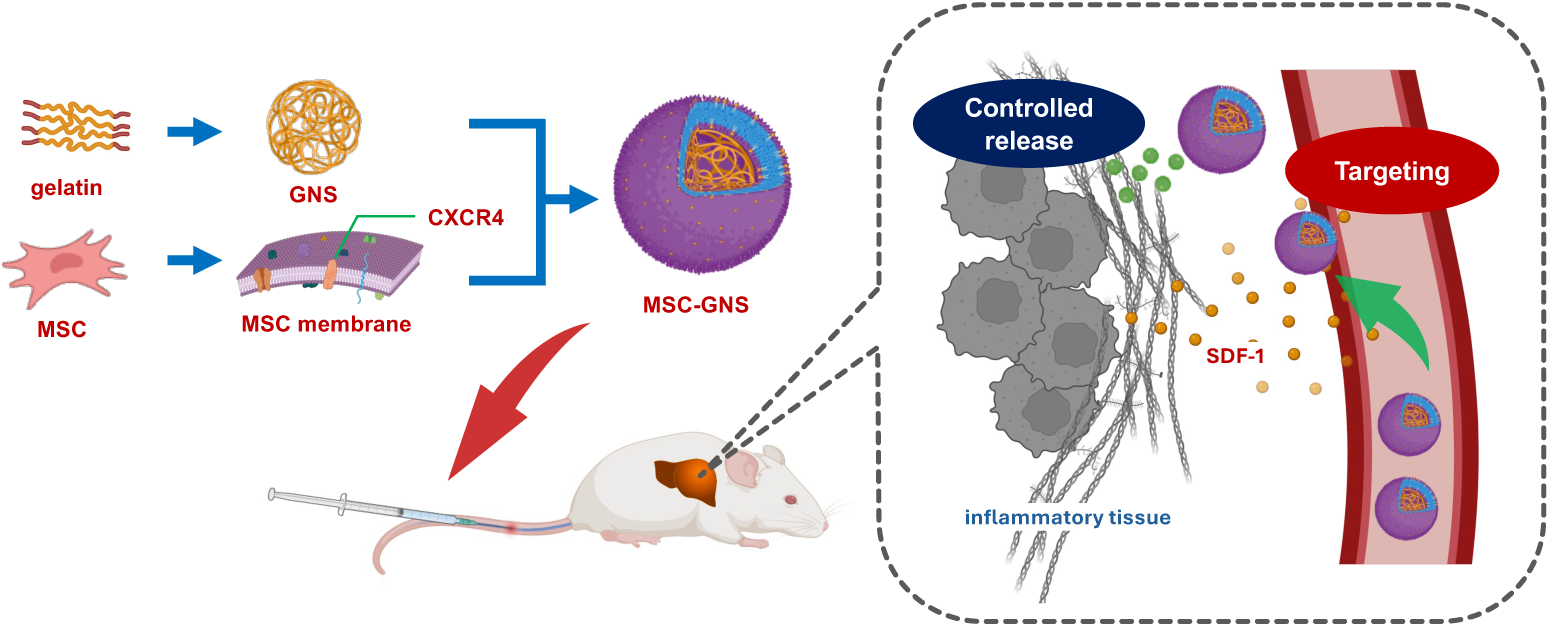

Mesenchymal stem cell membrane coated gelatin nanosphere as DDS nanocarrier

---

The effective suppression of inflammation by drug therapy requires prolonged drug efficacy, improved drug stability, accelerated absorption, and targeting to the treatment site. Drug delivery system (DDS) technology is important to achieve these drug effects.^1^ Nanospheres have been widely used as DDS carriers for therapeutic, prophylactic, and diagnostic applications because of their pharmacokinetic advantages, such as stability, enhanced absorption, and tissue distribution of pharmaceuticals.^2^

Nanospheres can be fabricated from various materials including lipids, iron oxide, silica, and biocompatible polymers, such as polylactic-co-glycolic acid (PLGA) and poly caprolactone.^2, 3^ Gelatin, a biocompatible protein derived from collagen, is enzymatically degradable, and used in pharmaceuticals, medical scaffolds, and food products.^4^ In our previous study, cationic gelatin was prepared by conjugating polyamines and gelatin conjugated with cholesteryl groups.^5^ Gelatin can be readily fabricated into various shapes, such as nanospheres,^6, 7^ microspheres,^8, 9^ films,^10^ and sponges.^11, 12^ In addition, gelatin nanospheres (GNS) are internalized into cells, and gradually release incorporated drugs, such as small interfering RNA (siRNA),^7^ molecular beacons,^6, 13^ and plasmid DNA^14^ in a degradation-dependent manner. Although an advantage of GNS is the controlled release of a drug, their tissue targeting ability remains to be improved. To regulate or suppress inflammation, it is necessary to allow GNS incorporating drugs to reach inflammatory tissues efficiently. The enhanced efficiency of drug delivery with minimal side effects should improve the tissue distribution of GNS.

Mesenchymal stem cells (MSC) have great potential for cell transplantation therapy. They have an inherent ability to accumulate in tumor and inflammatory tissues. One accumulation mechanism is the interaction between CXC-chemokine receptor 4 (CXCR4), a representative membrane protein expressed on MSC, and CXC motif chemokine ligand 12, also known as stromal cell-derived factor 1 (SDF-1).^15, 16^ Because SDF-1 is abundantly produced in inflammatory tissues, MSC are recruited to inflammatory tissues via a concentration-gradient mechanism.^17–19^ Various MSC membrane-coated nanoparticles have been reported,^20–22^ and these were shown to accumulate in large amounts in tumor, inflammatory, and ischemic sites.^23–25^

This study prepared MSC membrane-coated gelatin nanospheres (MSC-GNS) by coating GNS with MSC membranes, in order to enhance their tropism toward inflammatory tissues and achieve effective targeting and therapy for liver fibrosis. The morphology of MSC-GNS was observed, and western blotting and dot blotting analyses were performed to evaluate the coating. The drug release behavior and cellular uptake of MSC-GNS were examined. Finally, the tissue distribution of MSC-GNS after their intravenous injection into a mouse model of liver fibrosis and the subsequent therapeutic effect on liver fibrosis were evaluated.

## Results and Discussion

### Characterization of MSC-GNS

Various strategies can be used to provide nanospheres with tissue targeting abilities, such as the chemical modification of the nanosphere surface with antibodies, aptamers, or saccharides. The tissue distribution of nanospheres can be regulated by exploiting size differences or utilizing transporter activities that are dependent on pH.^26–30^ Recently, research in cell membrane-coated nanoparticles has increased.^31, 32^ Cell membranes have the advantages of biocompatibility and retaining various functions derived from the original cell type.^33^ Thus, the surface of nanospheres coated with cell membranes is expected to retain the functions of the original cell membranes. This technique will more easily mimic higher biological functions than the chemical modification of nanosphere surfaces. For example, red blood cell membrane-coated nanoparticles had improved blood retention,^34^ platelet membrane-coated nanoparticles specifically targeted a myocardial infarction site,^35^ and high numbers of cancer cell membrane-coated nanoparticles accumulated in tumors.^36^ In this study, a strategy of coating nanospheres with cell membranes was adopted to improve the targeting tropism. This technology enables the nanospheres to closely reconstitute the original functions of the donor cell membranes.^20, 31^ Another advantage of this strategy is to camouflage the nanospheres, protecting them from immune surveillance *in vivo*.

GNS were prepared by the conventional coacervation method with slight modification.^6^ The size of GNS can be changed by the molecular weight and concentration of the gelatin and the speed of stirring.^37^ Cell membrane-coated nanospheres have been prepared using various methods, such as extrusion, ultrasonication, and electroporation in microfluidic media.^31, 38, 39^ Thus, in this study, MSC-GNS were prepared using the extrusion and ultrasonication methods, and the results were compared to optimize the preparation method. **Table 1** shows the hydrodynamic size, polydispersity index, and ζ-potential of GNS, MSC-GNS, and MSC vesicles. The hydrodynamic size of GNS was 184.3 ± 3.1 nm, whereas that of MSC-GNS was slightly larger at 195.7 ± 5.5 nm. The ζ-potential of GNS was −3.3 ± 0.9 mV, whereas that of MSC-GNS was smaller at −10.7 ± 1.6 mV, which was similar to that of MSC vesicles (−11.4 ± 0.8 mV). Similar results have been reported by other researchers.^40, 41^ Taken together, the MSC membranes were shown to be successfully coated on the surface of GNS. Cationized gelatin nanospheres (cGNS) were prepared from ethylenediamine-conjugated gelatin with a size of 164.7 ± 3.4 nm and ζ-potential of 5.7 ± 0.7 mV. When attempting to coat cGNS with MSC membranes, they formed aggregates, which prevented their successful coating (**Table S1** and **Figure S1**). This phenomenon has also been reported by other studies.^42, 43^ It is possible that the balance of electrostatic forces between the cell membranes and nanoparticles was critical for successful cell membrane coating. **Figure 1A** and **1B** show the transmission electron microscope (TEM) and cryo-electron microscope (Cryo-EM) images of MSC-GNS and GNS. TEM observations showed that the entire particle was stained for GNS, whereas only the periphery was stained for MSC-GNS. This difference might be explained by the limited penetration of the staining agent into the interior of the particle in MSC-GNS. MSC-GNS and GNS had a spherical form. Compared with GNS, the peripheral area of MSC-GNS was dark by Cryo-EM observation. This finding has been reported for other cell membrane-coated nanospheres and liposomes^44, 45^ and may result from the difference in electron beam transmission, because of the phospholipids in the cell membrane component. **Figure 1C** shows the results of the single-particle analysis of MSC-GNS by imaging flow cytometry. The fluorescence signals of MSC_Rho_-GNS_QD705_ were scattered between the positive regions of GNS_QD705_ and MSC_Rho_ vesicles. Considering the cryo-EM observations, these findings strongly support the idea that MSC membranes covered the surface of GNS. The coating efficiency of MSC_Rho_-GNS_QD705_ prepared by the ultrasonication method was 94.1% but only 84.9% by the extrusion method. On the basis of this finding, the ultrasonication method was selected for the following experiments.

**Figure 1.**
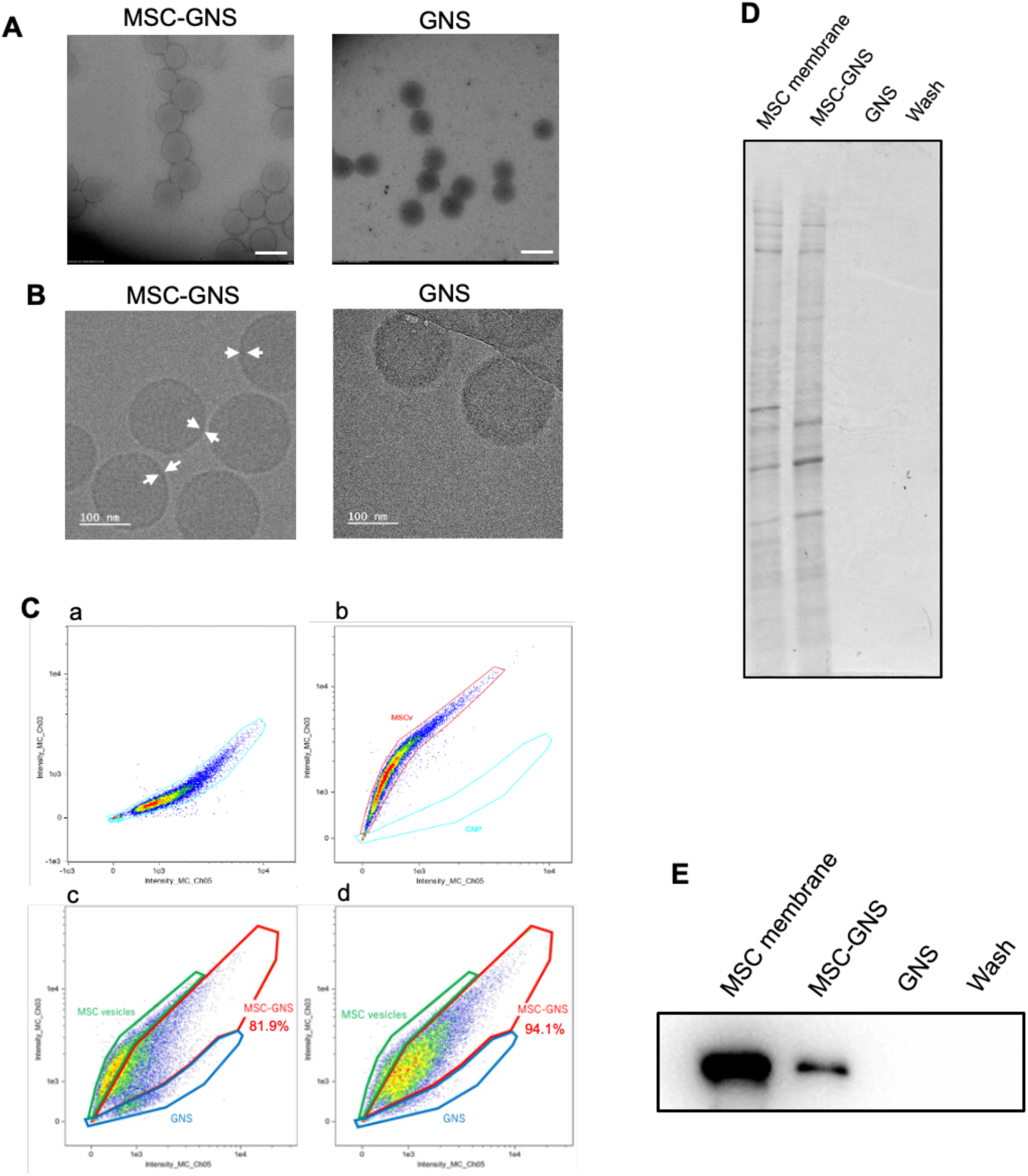
Characterization of MSC-GNS and GNS. (A) TEM images of MSC-GNS (left) and GNS (right). The scale bar is 200 nm. (B) Cryo-EM images of MSC-GNS (left) and GNS (right). The scale bar is 100 nm. The arrows indicate the dense layer localized on the GNS surface. (C) Imaging flow cytometry patterns of GNS (a), MSC membrane vesicles (b), MSC-GNS prepared by the extrusion method (c), and MSC-GNS prepared by the ultrasonication method (d). (D) SDS-PAGE profile of MSC membranes, MSC-GNS, GNS, and Wash sample. (E) Western blotting pattern of CXCR4 in MSC membranes, MSC-GNS, GNS, and Wash sample.

**Table 1.** Hydrodynamic size and ζ-potential of MSC-GNS and GNS.

| | Hydrodynamic size (nm) | PDI | ζ-potential (mV) |
| --- | --- | --- | --- |
| MSC-GNS | $195.7 \pm 5.5^a$ | 0.108 | $-10.7 \pm 1.6^a$ |
| GNS | $184.3 \pm 3.1$ | 0.053 | $-3.7 \pm 0.9$ |
| MSC vesicles | $264.3 \pm 22.4$ | 0.205 | $-11.4 \pm 0.8$ |

a): mean ± SD

### Protein analysis of isolated MSC membranes and MSC-GNS

**Figure S2A** and **S2B** show the western blotting results of isolated MSC membranes and MSC lysates. CXCR4 in the plasma membrane protein was more abundant in the isolated MSC membranes than in the cell lysates, whereas the cytoplasmic protein GAPDH was not detected. This indicated that cell membranes were successfully separated from cytoplasmic components. Other studies have reported isolation by ultrasonic methods using homogenizers or techniques with hypotonic lysates. The ultrasonic irradiation generates significant heat that has physical or chemical effects on the cell membranes structure and protein conformations, resulting in the loss of biological activity. Figure 1D shows the results of CBB staining. The protein band pattern of MSC-GNS was similar to that of MSC membranes, whereas no such pattern was observed for the GNS and Wash samples (coated operation with MSC membrane only, washed by centrifugation). These findings suggest that MSC membrane-derived proteins are retained on the surface of MSC-GNS. Figures 1E, **S3A,** and **S3B** show the results of CXCR4 detection by western blotting. Signals derived from CXCR4 were detected for MSC-GNS, but not for GNS or Wash samples. This indicated that excess MSC membrane vesicles were completely removed by centrifugation. These results were similar to those previously reported for other types of MSC nanoparticles.^23^ Next, the mixing ratio of MSC membranes to nanospheres was optimized for the preparation of MSC-GNS. Among the different mixing ratios, the amount of detected CXCR4 increased in an MSC membrane vesicle-dependent manner up to a 3:1 ratio, but after that remained nearly constant. This suggested that mixing GNS with MSC membranes at a 3:1 volume ratio was sufficient to coat GNS with MSC membranes. On the basis of these results, this ratio was used for the following experiments. However, the optimization of the mixing ratio requires careful consideration because it may vary depending on the evaluation method, coating technique, and physical properties and sizes of the core nanospheres. Indeed, Yang *et al.* determined the optimal mixing ratio of cell membranes to nanoparticles to be 1:5 by measuring the ζ-potential of MSC membrane-coated nanoparticles.^41^

### Membrane orientation of MSC-GNS and binding affinity of MSC-GNS for SDF-1

Although GNS can be coated with MSC cell membranes, CXCR4 cannot exert its biological functions unless the cell membranes are oriented in the same manner as their MSC origin. Therefore, the orientation of coated MSC membranes was investigated by detecting the ectodomain of intact CXCR4 under non-denaturing conditions by dot blotting. It has been reported that the orientation of a protein can be determined by detecting each protein domain using antibodies with known epitopes. This study used an antibody that recognizes the ectodomain of CXCR4; therefore, if the cell membrane was coated in the same orientation as the MSC origin, signals derived from CXCR4 can be detected. Figure 2A shows the dot blotting results. Dots of CXCR4 were detected in the MSC-GNS group, but not the GNS group. This clearly indicates that the ectodomain of CXCR4 protruded from MSC-GNS. It is likely that the orientation of MSC membranes on MSC-GNS is retained similar to that of the original MSC membranes. MSC exhibit chemotaxis toward inflammatory tissues through the binding of CXCR4 on MSC to the chemokine SDF-1, which is secreted predominantly in inflammatory tissues.^18, 46–48^ Therefore, given the importance of the binding of MSC-GNS to SDF-1 in targeting inflammatory tissues, we examined whether MSC-GNS binds to SDF-1. Figure 2B shows the evaluation results of the binding ability of MSC-GNS to SDF-1. SDF-1 was detected strongly in the MSC-GNS group, indicating that CXCR4 on the MSC-GNS maintained its binding ability for SDF-1. Although a weak signal was also detected in the GNS group, this might be explained by the nonspecific adsorption of SDF-1 by the GNS.^49^ To the best of our knowledge, this is the first direct evaluation of a specific interaction on cell membrane-coated nanospheres.

**Figure 2.**
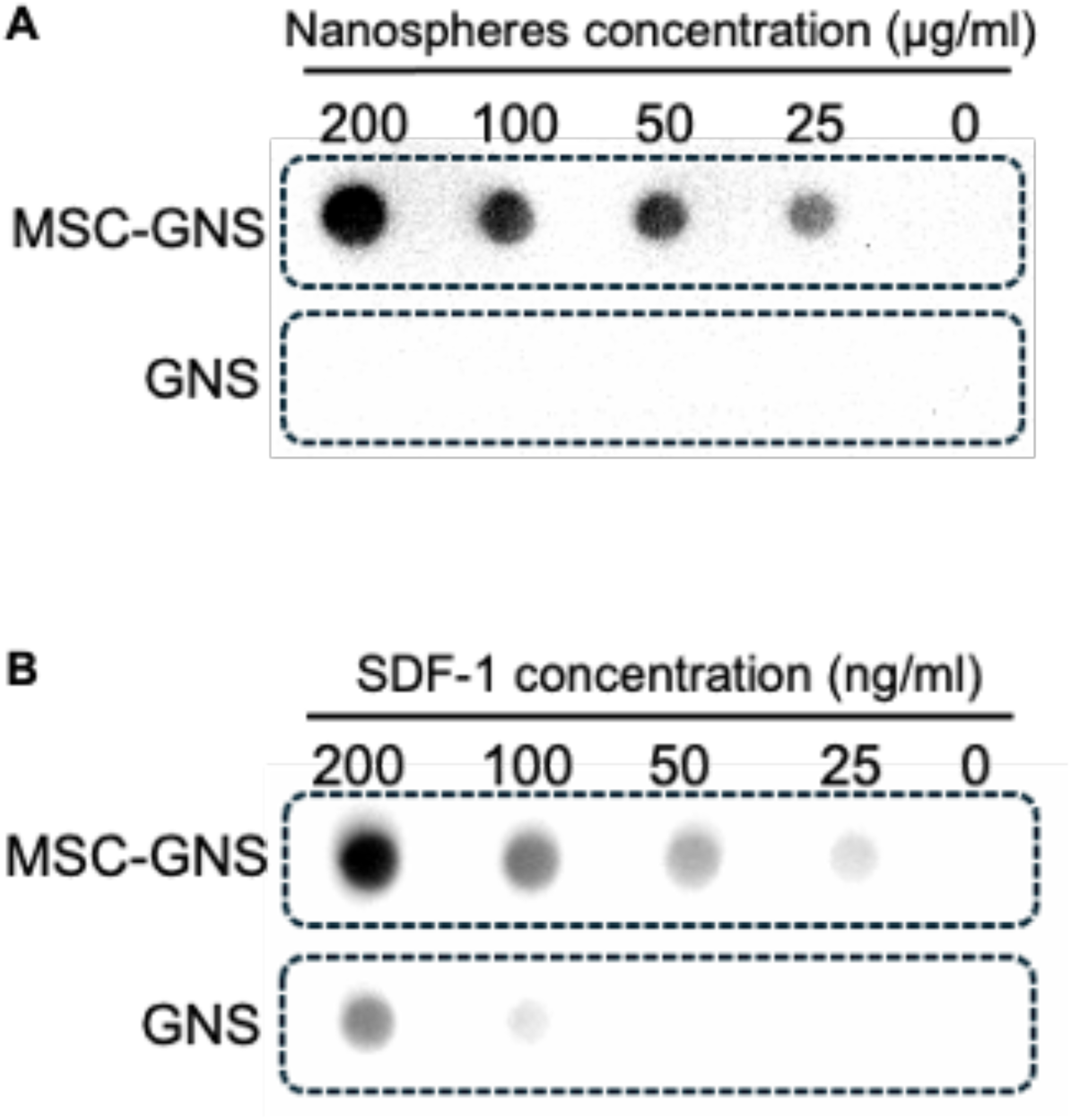
Membrane orientation of MSC-GNS and binding affinity of MSC-GNS for SDF-1. (A) Dot blotting pattern of the ectodomain of CXCR4 in MSC-GNS and GNS at different nanosphere concentrations. (B) Dot blotting pattern of SDF1 bound to MSC-GNS and GNS after incubation with different SDF-1 concentrations.

### Collagenase-dependent degradation and controlled drug release from MSC-GNS

MSC membranes were successfully coated onto GNS without losing their biological function. Next, we verified whether coating GNS with MSC membranes impaired their intrinsic controlled drug-releasing ability. Figure 3A shows a time-course degradation profile of GNS and MSC-GNS in the presence or absence of collagenase. Little degradation was observed for MSC-GNS and GNS in the absence of collagenase. In contrast, in the presence of collagenase, both were gradually degraded over time. MSC-GNS were degraded more slowly than GNS. High performance liquid chromatography (HPLC) result indicated that LSKL was successfully encapsulated into GNS with an incorporation efficiency of 63.7 ± 2.0% (**Figure S4**). Figure 3B shows the release profile of LSKL from MSC-GNS and GNS in the presence or absence of collagenase. In the presence of collagenase, LSKL was released from MSC-GNS and GNS, corresponding to their degradation behavior described above. This suggests that LSKL is released as GNS degrades. Similar to the degradation behavior of GNS, LSKL release was delayed in MSC-GNS compared with GNS. A previous study reported that drug release was delayed in macrophage membrane-coated PLGA nanoparticles compared with uncoated nanoparticles.^50^ This may be because the cell membrane on nanoparticles acts as an interface, preventing collagenase from mediating their degradation. Furthermore, with prolonged incubation with a high concentration of collagenase under refined conditions, the protein components on the surface of the MSC-GNS tended to degrade nonspecifically, leading to the subsequent degradation of the internal GNS. In previous studies, the degradation kinetics of GNS were controlled by regulating the degree of cross-linking inside the GNS, thereby influencing the release rate of the incorporated drug.^7^ Thus, even in the case of MSC-GNS, it may be possible to regulate artificially the release behavior of the encapsulated drug by altering cross-linking within the GNS.

**Figure 3.**
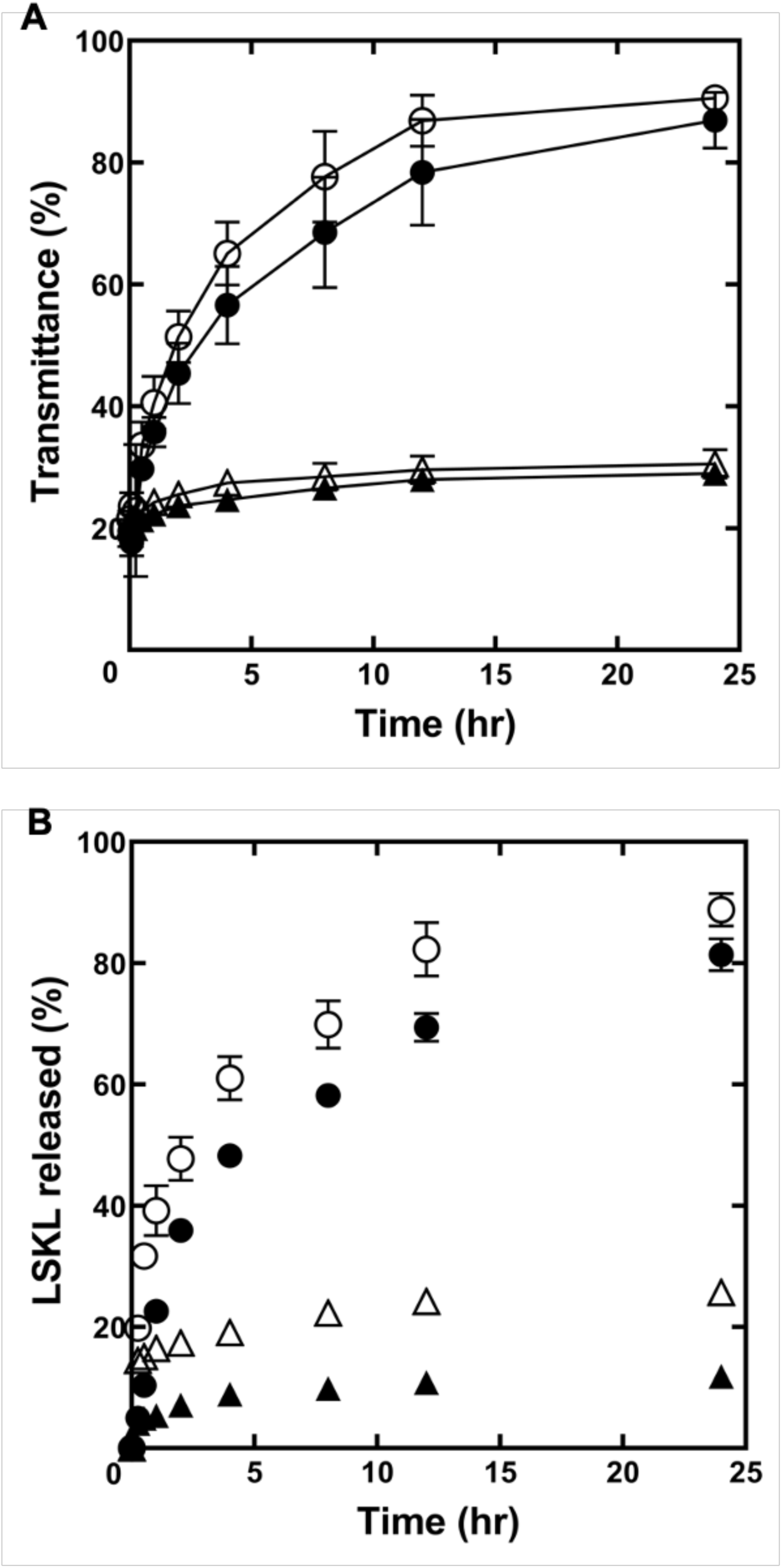
Collagenase-dependent degradation and controlled drug release from MSC-GNS. (A) Time profiles of MSC-GNS and GNS degradation in PBS with (circle) or without collagenase (triangle). (B) Time profiles of LSKL release from MSC-GNS and GNS in PBS with or without collagenase: MSC-GNS (●, ▴) and GNS (○, △). All results represent the average ± SD (*n* = 3).

### Blood clearance and cellular uptake of MSC-GNS

Figure 4A shows the time course of the blood concentration of nanospheres after their intravenous injection into mice. Both types of nanospheres exhibited biphasic blood retention; therefore, they were analyzed using a two-compartment model (**Table 2**). During the early distribution phase, there was no difference in the blood retention of the nanospheres. However, during the later elimination phase, MSC-GNS had a higher retention than GNS, with its half-life approximately doubling in this phase. This result was similar to that observed for erythrocyte membrane-coated nanoparticles.^44^ In general, during the elimination phase, the drug concentration is considered to decrease only through the process of elimination, because the drug distribution between major and peripheral tissues has reached equilibrium.^51^ At this time, the drug is mainly eliminated *in vivo* by immune cells. It is well known that hematopoietic cells evade attacks from immune cells, and that this function occurs in the cell membrane.^52^ To investigate the cause of the difference in the elimination phase, the uptake of each nanosphere into cells was evaluated. Incubation with MSC-GNS or GNS did not induce cytotoxicity against RAW264.7 cells or MC3T3-E1 cells at various concentrations (**Figure S5**). This result differs from other types of MSC membrane-coated nanospheres, which are cytotoxic above a certain concentration.^41, 53, 54^ Figure 4B shows confocal laser scanning microscopy images of RAW264.7 cells and MC3T3-E1 cells after incubation with GNS or MSC-GNS. Fluorescent signals derived from the Qdot 605 internalized into GNS were observed in both cell lines. There was no difference in the fluorescence signals between the GNS and MSC-GNS groups in MC3T3-E1 cells. However, in RAW264.7 cells, the fluorescence signals in the MSC-GNS group were lower than those in the GNS. Figure 4C and **4D** show the quantitative results of the cellular uptake of nanospheres by flow cytometry. The results were consistent with confocal laser microscopic observations, showing that MSC-GNS was significantly less incorporated in RAW264.7 cells than GNS. Unlike the results for RAW264.7 cells, the uptake of MSC-GNS into MC3T3-E1 cells was comparable to that of GNS. This suggests that the cell membrane coating did not affect the uptake into non-immune cells. The MSC membranes on the surface of GNS would be less likely to be taken up by macrophage-like cells. Therefore, MSC-GNS may evade phagocytosis by macrophages *in vivo*, leading to an extended blood retention time. Bose *et al*. fabricated MSC-coated PLGA nanospheres that had a reduced uptake into macrophage-like J774 and THP cells.^23^ Similarly, Wang *et al*. prepared red blood cell membrane-coated PLGA nanospheres, which exhibited reduced uptake into RAW264.7 cells.^55^ These findings can be explained by the function of CD47, a membrane protein on the plasma membrane surface of hematopoietic cells. It is well known that CD47 functions as a “don’t eat me signal”, which inhibits macrophage phagocytosis when it binds to SIRPα, a membrane protein on the surface of macrophages.^52, 56, 57^ Indeed, CD47 was detected in MSC-GNS by western blotting (**Figure S6**). Therefore, it is highly likely that CD47 on MSC-GNS inhibits phagocytosis, leading to its delayed elimination from the blood and extended retention time during the late elimination phase. A previous study showed that MSC membrane-coating increased the uptake into fibroblasts and vascular endothelial cells.^58^ However, in this study, the uptake into MC3T3-E1 cells was not significant. However, our study used different types of nanoparticles and different coating methods for MSC membranes. Therefore, more detailed studies are needed to determine the changes in cellular uptake by non-immune cells.

**Figure 4.**
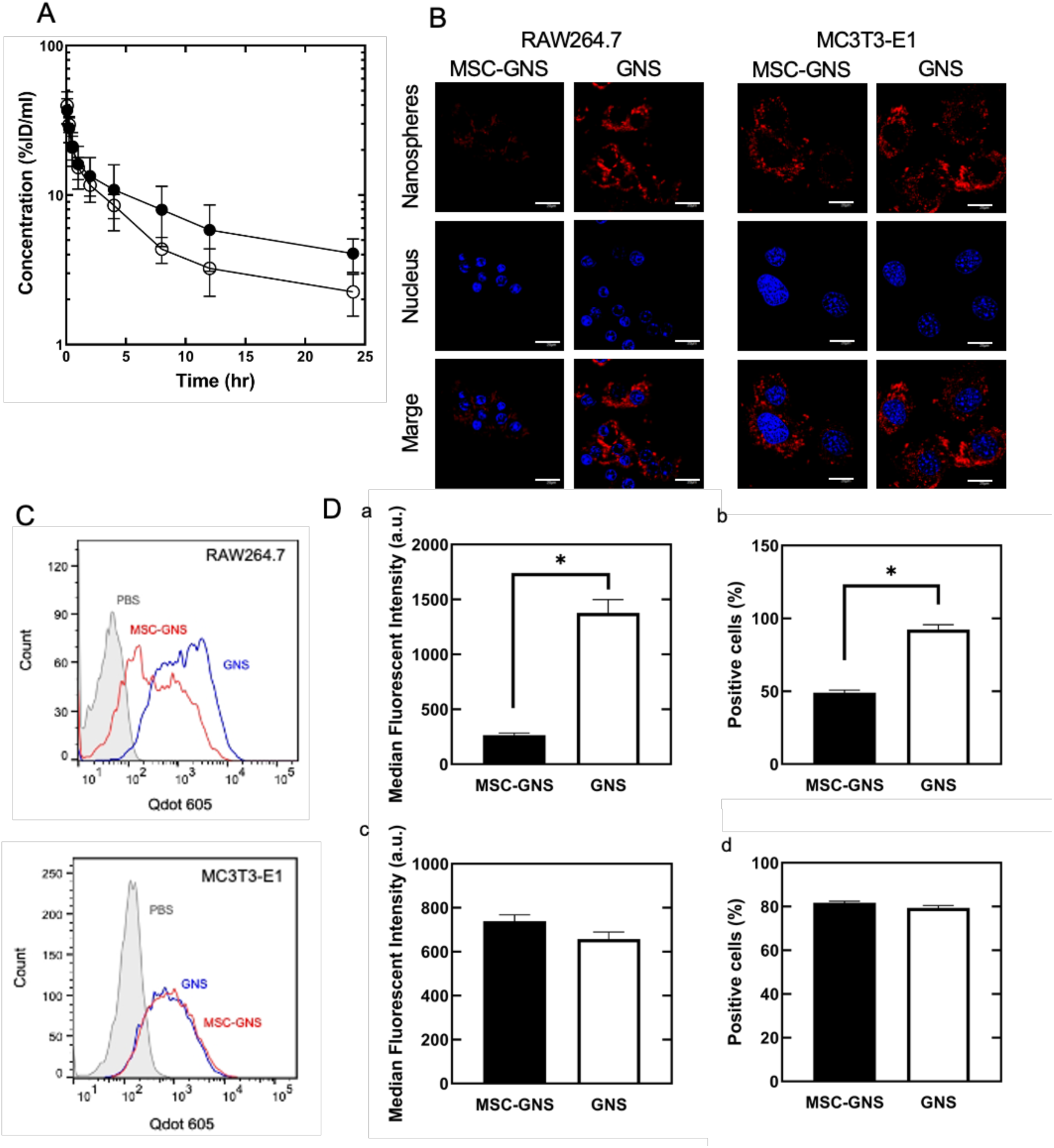
Blood clearance and cellular uptake of MSC-GNS. (A) Time profiles of MSC-GNS and GNS in the circulating blood 24 hours after intravenous injection. (B) Confocal laser scanning microscopic images of RAW264.7 and MC3T3-E1 cells 6 and 12 hours after incubation with MSC-GNS or GNS, respectively. The scale bar is 20 μm. MSC-GNS and GNS incorporate Qdot605. Cell nuclei were stained by Hoechst 33342. (C) Single-cell analysis of internalized MSC-GNS and GNS in RAW264.7 (top) and MC3T3-E1 (bottom) cells 6 and 12 hours after incubation with MSC-GNS and GNS, respectively. (D) Cell uptake of MSC-GNS and GNS by RAW264.7 (a, b) and MC3T3-E1 (c, d) cells. (a, c) Median fluorescence intensity of nanospheres. (b, d) The percentage of nanospheres taken up by cells. All results represent the average ± SD (n = 3). *p < 0.05, two-tailed Welch’s *t*-test.

**Table 2.** Pharmacokinetic and pharmacodynamic values of MSC-GNS and GNS.

|  | AUC<br>(hr · µg/ml) | MRT<br>(hr) | CL<br>(ml/hr) | V<sub>ss</sub><br>(ml) | t<sub>1/2(α)</sub><br>(hr) | t<sub>1/2(β)</sub><br>(hr) |
| --- | --- | --- | --- | --- | --- | --- |
| MSC-GNS | 178±57.8<sup>a)</sup> | 26.2±23.9 | 0.561±0.486 | 14.7±13.0 | 0.242 | 9.92 |
| GNS | 109±33.4 | 20.0±13.8 | 0.916±0.281 | 18.4±7.51 | 0.232 | 5.10 |

a): mean ± SD

### *Ex vivo* imaging of nanospheres

Liver fibrosis, a chronic liver disease, is a global health challenge. As liver fibrosis progresses, it can lead to irreversible cirrhosis and liver cancer, and the early treatment of liver fibrosis is needed.^59^ A liver fibrosis model was established by the intraperitoneal injection of carbon tetrachloride into mice. To confirm the establishment of the model, we performed hematoxylin and eosin (HE) staining and Masson’s trichrome (MT) staining of livers (**Figure S7A**), and plasma concentrations of alanine aminotransferase (ALT) and aspartate aminotransferase (AST) were measured 4 weeks after the first injection of carbon tetrachloride (**Figure S7B**). As a result, the plasma concentrations of ALT and AST were significantly increased, whereas fibrotic areas were observed at the end of the experiment. Furthermore, the concentration of SDF-1 in livers and plasma was significantly increased in mice treated with carbon tetrachloride (**Figure S7C**), as reported in previous studies.^60^ Figure 5A and **5B** show the tissue distribution of nanospheres 3 and 24 hours after intravenous injection. Most nanospheres had accumulated in the liver, irrespective of the presence of liver fibrosis. Importantly, although the accumulation of MSC-GNS or GNS in the normal liver was almost comparable, the accumulation of MSC-GNS was ∼ 1.7 times higher than that of GNS in fibrotic livers. MSC migrate to fibrotic livers *via* the CXCR4-SDF-1 axis.^61^ Furthermore, in the liver, hepatic astrocytes activated by fibrosis and other factors secrete SDF-1 to mobilize CXCR4-positive cells, further promoting α-smooth muscle actin (α-SMA) production by the surrounding myofibroblasts.^62^ Therefore, GNS coated with CXCR4-positive MSC membranes accumulated in the fibrotic livers. Previous studies reported that MSC membrane-coated nanoparticles accumulated in inflammatory lesions related to cancer,^63^ as well as ischemic limbs,^23^ and myocardial infarction hearts.^24^ Overall, it is difficult to explain the liver accumulation of MSC membrane-coated nanoparticles in the liver fibrosis model using only one mechanism; however, their selectivity for inflammatory tissues may have had a positive effect.

**Figure 5.**
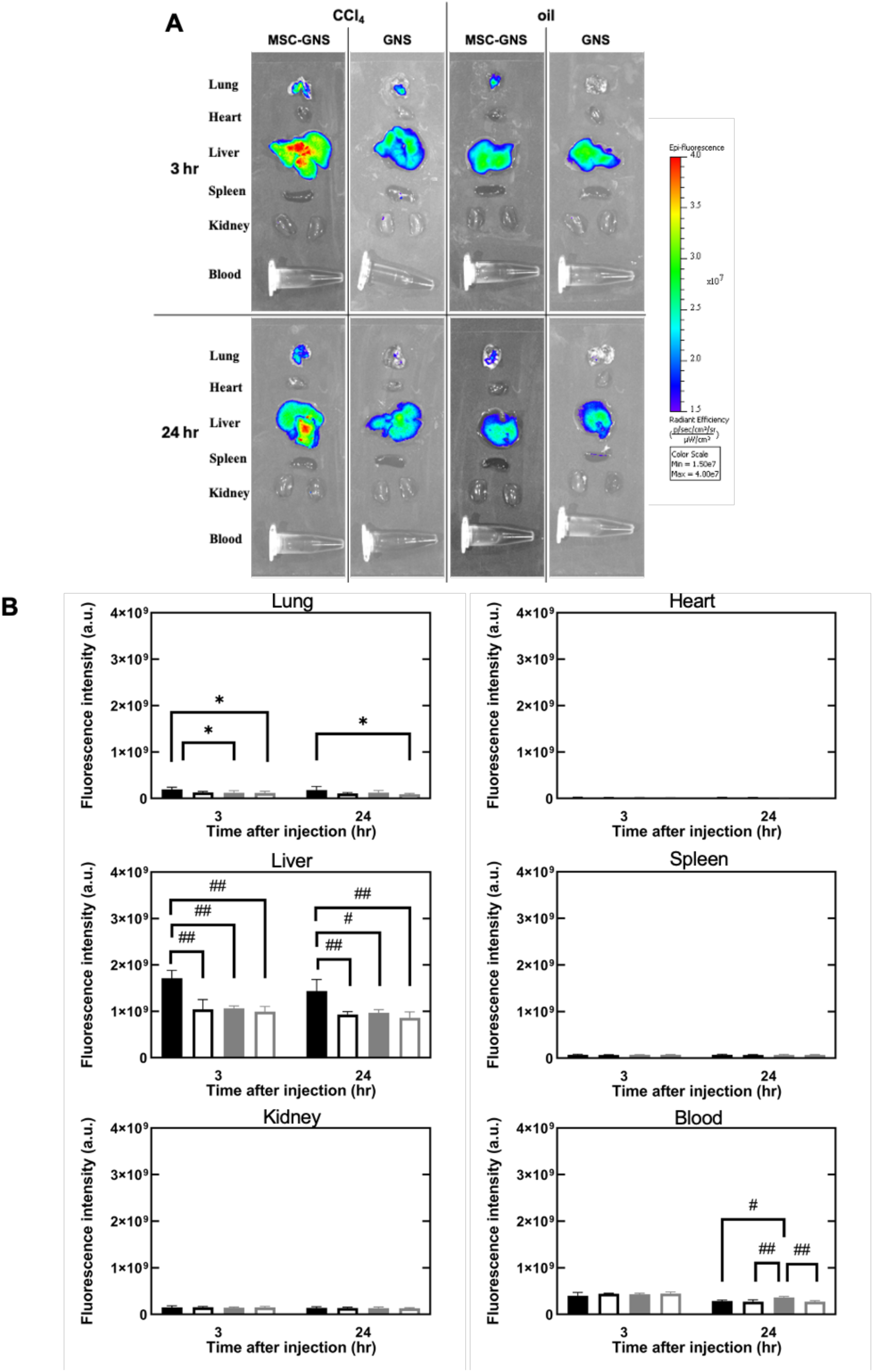
**Tissue-distribution of MSC-GNS. (**A) Tissue-distribution images of MSC-GNS and GNS in liver fibrotic (CCl4-treated) and normal (olive oil-treated) mice 3 and 24 hours after intravenous injection. (B) Quantitative accumulation patterns of MSC-GNS and GNS 24 hours after intravenous injection: MSC-GNS injected to liver fibrosis model mice (black closed square), GNS injected to liver fibrosis model mice (black open square), MSC-GNS injected to normal mice (gray closed square), and MSC-GNS injected to l liver fibrosis model mice (gray open square). The data are presented as the mean ± s.d. (*n* = 6) One-way factorial ANOVA: *Adjusted p < 0.05, **: p < 0.01, #: p < 0.005, and ##: p < 0.0001, Tukey’s post hoc test.

### Therapeutic effect on liver fibrosis

LSKL is a 4-amino acid peptide composed of leucine, serine, lysine, and leucine. LSKL competitively inhibits thrombospondin (TSP-1), which prevents LAP-TGF-β1 from maturating active TGF-β1, resulting in the inhibition of fibrosis progression.^64, 65^ Previous studies reported the injection of LSKL suppressed chronic tissue inflammation including liver fibrosis,^66^ renal interstitial fibrosis,^67^ and subarachnoid fibrosis.^68^ In the current study, MSC-GNS-LSKL, GNS-LSKL, and LSKL were injected via the tail vein into liver fibrosis model mice for 2 weeks, for a total of six doses (Figure 6A). Figure 6B shows the histological observations after MSC-GNS-LSKL injection by HE and MT liver staining. Figure 6C shows the percentage of fibrotic areas observed. MT staining demonstrated the injection of GNS-LSKL and MSC-GNS-LSKL led to a reduction in fibrotic areas, whereas the injection of LSKL alone did not reduce fibrotic areas compared with saline. The fibrotic area was decreased in the following order: LSKL, GNS-LSKL, and MSC-GNS-LSKL. Figure 6D **and 6E** show the plasma concentrations of ALT and AST after treatment. The ALT and AST concentrations in mice injected with LSKL alone were not significantly changed compared with those in saline-injected mice. The MSC-GNS-LSKL group had the lowest ALT and AST concentrations of all groups. This indicated that MSC-GNS-LSKL treatment reduced liver damage and the progression to liver fibrosis. Next, the expression levels of α-SMA and β-actin in livers were evaluated (Figure 6F), because the expression of α-SMA increases with fibrosis.^69^ The MSC-GNS-LSKL injected groups had the highest reduction in the expression levels of α-SMA. Taken together with the histological observations and α-SMA expression, the therapeutic effect might have been achieved by targeting and controlling the release of LSKL using MSC-GNS. In addition, MSC-GNS-LSKL had a therapeutic effect demonstrated by the suppression of liver fibrosis at a lower dose compared with the intraperitoneal injection of LSKL alone in a previous study.^66^ The MSC membrane-coated nanoparticles also had higher therapeutic efficacy than uncoated nanoparticles in inflammatory diseases other than liver fibrosis, such as critical limb ischemia and myocardial infarction.^23, 24^ Therefore, MSC membrane-coated nanoparticles may be applicable for the treatment of a wide range of inflammatory diseases.

**Figure 6.**
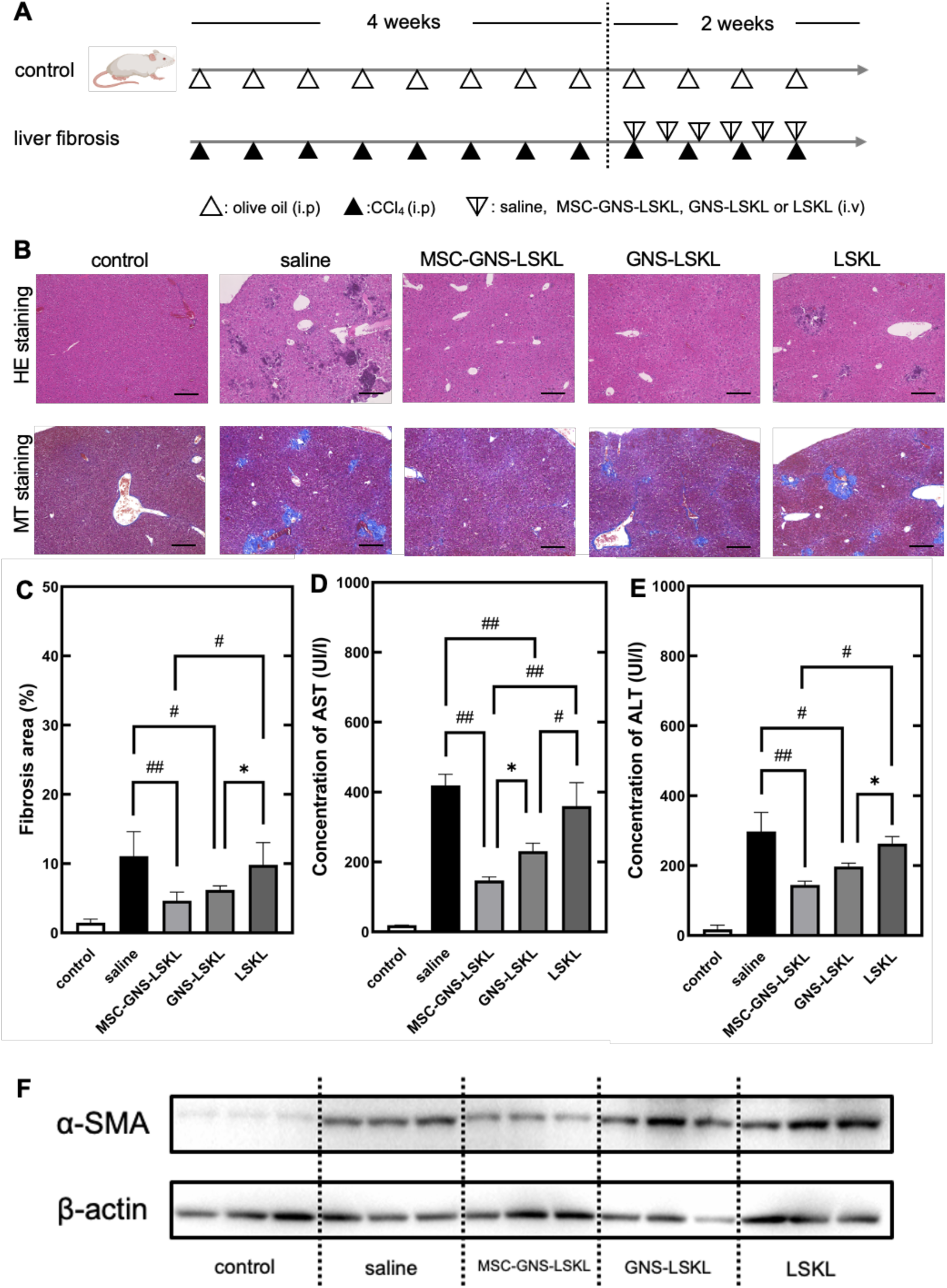
Therapeutic effect of MSC-GNS. (A) Therapeutic injection schedule of MSC-GNS-LSKL, GNS-LSKL, and LSKL. (B) HE and MT staining images of mouse livers two days after the last injection. The scale bar is 200 μm. (C) Liver fibrotic areas by MT staining. The data are presented as the mean ± s.d. (*n* = 3). Plasma concentrations of (D) AST and (E) ALT in treated mice two days after final injection of therapeutics. The data are presented as the mean ± s.d. (*n* = 6). (F) Western blotting of α-SMA and β-actin in the liver of mice treated. One-way factorial ANOVA: *Adjusted p < 0.05, **: p < 0.01, #: p < 0.005, and ##: p < 0.0001, Tukey’s post hoc test.

## Conclusion

In this study, MSC-GNS were prepared by coating GNS, a drug release carrier, with MSC membranes. The effects of the coating were evaluated by cryo-electron microscopy, single particle analysis, western blotting, and dot blotting. One characteristic of MSC-GNS is controlled drug release, even when coated with cell membranes. Following their intravenous injection into a mouse model of liver fibrosis, MSC-GNS exhibited greater accumulation in inflamed livers and higher blood retention compared with MSC-membrane-free GNS. Furthermore, MSC-GNS loaded with anti-fibrotic agents had an enhanced therapeutic effect on liver fibrosis. Therefore, MSC-GNS is a drug carrier with inflammatory tissue targeting and controlled drug release abilities.

## Materials and Methods

### Cell culture

The murine bone marrow-derived mesenchymal stem cell line KUM6 was obtained from the Japanese Collection of Research Bioresources Cell Bank (JCRB) (Suita, Japan). KUM6 cells were cultured in Iscove’s Modified Dulbecco’s Medium (IMDM; Thermo Fisher Scientific Inc., Waltham, MA, USA) supplemented with 10 vol% fetal calf serum (FCS) and 1 vol% penicillin/streptomycin at 37 °C in an incubator with 5% CO_2_. The murine leukemia macrophage cell line RAW264.7 was obtained from RIKEN BioResource Research Center (RIKEN BRC) (Tsukuba, Japan) and cultured in Roswell Park Memorial Institute medium-1640 (RPMI 1640, Thermo Fisher Scientific Inc.) with 10 vol% FCS and 1 vol% penicillin/streptomycin at 37 °C in 5% CO_2_. The murine osteoblast precursor cell line MC3T3-E1 was cultured in minimal essential medium α (MEM-α, Thermo Fisher Scientific Inc.) with 10 vol% FCS and 1 vol% penicillin/streptomycin at 37 °C in 5% CO_2_.^70^

### Isolation of MSC membranes

When KUM6 cell confluence reached 90%, cells were washed in phosphate-buffered saline (PBS, pH 7.4) and detached by trypsin-ethylenediaminetetraacetic acid. After the cell suspension was centrifuged at 200 ×g for 3 minutes and washed in PBS three times, the cells were resuspended in PBS. The cell dispersion was frozen and re-thawed three times at −80 °C and room temperature, then centrifuged at 4000 ×g for 3 minutes four times and resuspended in double-distilled water (DDW). The isolated cell membrane was stored at −80 °C for subsequent experiments. The protein concentrations of cell membranes were quantified using a BCA protein assay kit (Thermo Fisher Scientific Inc.).

### Preparation of gelatin nanospheres (GNS)

Gelatin (isoelectric point: 9.0, weight-averaged molecular weight: 100,000) was kindly supplied by Nitta Gelatin Inc., Osaka, Japan. GNS were prepared by a slight modification of the conventional coacervation method.^6^ Briefly, 1.25 ml of gelatin aqueous solution (20.0 mg/ml) was prepared by warming at 40 °C. Next, 25 µl of 1 M HCl (Nacalai Tesque Inc., Kyoto, Japan) was added to the solution, and. 5.0 ml of acetone (Nacalai Tesque Inc.) was dropped at 1.0 ml/min through a 5 ml syringe with a 24-gauge needle into the solution with stirring at 1000 rpm to form the coacervate. Then, 20 µl of 25 wt % glutaraldehyde (Nacalai Tesque Inc.) was added to allow the chemical cross-linking of GNS for 6 hours. Next, 2.0 ml of aqueous glycine solution (37.5 mg/ml) was added to block the unreacted aldehyde groups. The resulting solution was stirred at room temperature overnight and acetone was evaporated. After the solution was centrifuged at 16,000 ×g for 30 minutes at 4 ℃, the GNS pellet was collected, resuspended in DDW, and centrifuged at 1000 ×g for 3 minutes. Then, the supernatant was centrifuged at 16,000 ×g for 30 minutes and washed in DDW twice. The concentration of GNS produced was determined using a Pierce BCA Protein Assay Kit (Thermo Fisher Scientific Inc.).

### Preparation of GNS incorporating LSKL (GNS-LSKL)

One mg of GNS was freeze-dried, and 20 µl of 50 mg/ml LSKL tetrapeptide solution (MedChemExpress Inc., Monmouth Junction, NJ, USA) was added to the freeze-dried GNS and allowed to stand for 15 minutes at 37 °C. Then, the GNS-LSKL solution was incubated at 37 °C for 3 hours with agitation, and freeze-dried again. After freeze-drying, samples were resuspended with 1 ml of PBS and centrifuged at 16,000 ×g for 30 minutes. The supernatant was collected, and the residue of LSKL was evaluated by HPLC (Prominence LC-20AT; Shimazu Co., Ltd., Kyoto, Japan). A C_18_ column was used for HPLC, and H_2_O containing 0.1% trifluoroacetic acid and acetonitrile (Nacalai Tesque Inc.) containing 0.1% trifluoroacetic acid (Nacalai Tesque Inc.) were used as mobile phases. The LSKL incorporating rate was calculated using the following equation.

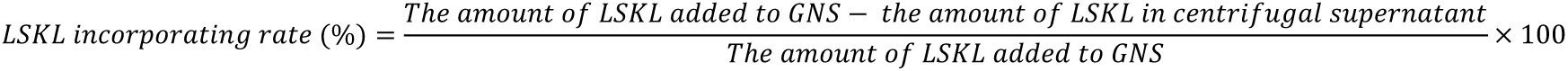

### Preparation of MSC membrane-coated gelatin nanospheres (MSC-GNS)

The coating of MSC membranes on the surface of GNS was performed by the extrusion or ultrasonication methods. Isolated MSC membranes were extruded 21 times through a polycarbonate membrane with a mesh size of 1 µm using a Mini-Extruder (Avanti Polar Lipids Inc., Alabaster, AL, USA) to prepare the MSC membrane vesicles. In the extrusion method, MSC membrane vesicles and GNS were mixed so that the protein weight ratio was 3:1 and the mixture was extruded 21 times through a polycarbonate membrane with a mesh size of 400 nm. For the ultrasonication method, MSC membrane vesicles were extruded 21 times through a polycarbonate membrane with a mesh size of 400 nm, mixed with GNS, and then ultrasonicated (55 W, 42 kHz) for 10 minutes. After coating, they were centrifuged at 14,000 ×g for 30 minutes and washed three times to remove excess MSC membrane vesicles. Finally, MSC-GNS were collected. MSC membrane-coated GNS-LSKL (MSC-GNS-LSKL) were prepared by coating GNS-LSKL with MSC membranes using the same method as above.

### Measurement of the hydrodynamic size and ζ-potential

The hydrodynamic size of GNS and MSC-GNS were measured by dynamic light scattering (DLS) using a Zetasizer Nano-ZS (Malvern Instruments Ltd., Worcestershire, UK).

Before the measurement, nanospheres were resuspended in 10 mM PBS. The ζ-potential of nanospheres was measured by electrophoresis light scattering (ELS) using a Zetasizer Nano-ZS. Before the measurement, nanospheres were resuspended in 10 mM phosphate buffer.

### Transmission electron microscopy

The morphology of the nanospheres was observed using a TEM (HT7700-TEM, Hitachi Corp., Japan), operated at 100 kV. First, 10 μl of nanosphere solution was deposited on a copper grid coated with an elastic carbon film (Elastic Wetties, Nissin EM Co., Ltd., Japan) for 30 minutes. After removing excess fluid by blotting, the grids were fixed with 4% paraformaldehyde (Pierce™ 16% Formaldehyde (w/v), Methanol-free, Thermo Fisher Scientific Inc.) for 15 minutes. Before observation by TEM, they were washed with DDW and negatively stained with EM stainer (Nissin EM Co., Ltd.) for 30 minutes.

### Cryo-electron microscopy

The morphology of the nanospheres was observed using a Cryo-EM (JEM-2100Plus, JEOL Co., Ltd., Japan), operated at 200 kV. First, 5 μl of nanosphere solution was deposited on holey carbon support film grids (Quantifoil R2/1, EM-Japan Co., Ltd., Japan) for 1 minute. After removing excess fluid by blotting, the grids were then directly plunged into liquid ethane. Grids were stored in liquid nitrogen until transferred to the electron microscope.

### Single particle analysis

MSC-GNS were analyzed for single particles using imaging flow cytometry (AMNIS ImageStream MarkⅡFlow Cytometer, Cytek Biosciences Inc., Fremont, CA, USA). MSC membranes were mixed with 1,2-dioleoyl-sn-glycero-3-phosphoethanolamine-N-(lissamine rhodamine B sulfonyl) (ammonium salt) (0.768 mM) (Avanti Polar Lipids Inc.) and incubated in the dark for 30 minutes to obtain rhodamine B labeled MSC membranes (MSC_Rho_). Qdots-encapsulated GNS (GNS_QD705_) were prepared by mixing a gelatin solution with Qdot705 (Qdot 705 ITK Carboxyl Quantum Dots, Thermo Fisher Scientific Inc.), which formed a coacervate. MSC_Rho_-GNS_QD705_ were prepared as described above. Signals were detected at Ch01 for brightfield, Ch03 for GNS_QD705_, Ch05 for MSC_Rho_, and Ch06 for side scattered light (SSC), respectively. The same gating strategy was used for all samples. Each sample contained 20,000 particles, and detection was focused on the MSC_Rho_ and GNS_QD705_ double-positive regions. Data analysis was performed using IDEAS software (Cytek Biosciences Inc.).

### Total protein analysis of MSC-GNS

The total protein concentrations of MSC-GNS and GNS were measured by BCA assay.

SDS-polyacrylamide gel electrophoresis (SDS-PAGE) of a 10% polyacrylamide gel (10% Mini-Protean TGX Gel, Bio-Rad Laboratories Inc., Hercules, CA, USA) was performed, and the gel was stained with Coomassie Brilliant Blue (Bullet CBB Stain One; Nacalai Tesque Inc.) at 4 °C overnight. After destaining, the gel was visualized using a Bio-Rad GelDoc Go system (Bio-Rad Laboratories Inc.).

### Western blot analysis of MSC-GNS

Samples were mixed with Laemmli buffer containing 10% 2-mercaptoethanol and incubated at 70 °C for 15 minutes. Then, they were separated by SDS-PAGE under reducing conditions. Subsequently, bands were electrotransferred to a polyvinylidene difluoride (PVDF) membrane (Trans-Blot Turbo 0.2-micron PVDF Membrane, Bio-Rad Laboratories Inc.) at 4 ℃ for 16 hours. The PVDF membrane was incubated with Blocking One (Nacalai Tesque Inc.) at room temperature for 1 hour and washed using Tris-buffered saline containing 0.02% Tween-20 (TBS-T) three times. Next, the membrane was incubated with anti-CXCR4 antibodies (4G10, 200 ng/ml; Santa Cruz Biotechnology, Inc., Santa Cruz, CA, USA) in TBS-T containing 10% Blocking One at 4 ℃ overnight. After washing with TBS-T, the membrane was incubated with anti-mouse IgG HRP-conjugated antibodies (20 ng/ml; Cell Signaling Technology Inc., Danvers, MA, USA) at room temperature for 1 hour. Then, after washing with TBS-T, the membrane was incubated with the electrochemical luminescence (ECL) substrate (Pierce ECL Plus Western Blotting Substrate, Thermo Fisher Scientific Inc.) and exposed using a Bio-Rad ChemiDoc XRS plus system (Bio-Rad Laboratories Inc.).

Liver samples were homogenized in lysis buffer, andα-SMA and β-actin in liver lysates were detected by western blotting. Briefly, SDS-PAGE was performed, and the bands were electrotransferred to a PVDF membrane, followed by incubation with Blocking One or Blocking One-P for phosphorylated proteins (Nacalai Tesque Inc.). Then, the membrane was incubated with anti-α-SMA antibodies (ab7817, 500 ng/ml; Abcam Ltd.) and anti-β-actin antibodies (#4970, 61 ng/ml; Cell Signaling Technology Inc.). After incubation with anti-mouse IgG HRP-conjugated antibodies and anti-rabbit IgG HRP-conjugated antibodies, a luminescent substrate was added before imaging.

### Evaluation of the orientation of MSC membranes on GNS

MSC-GNS and GNS were incubated on nitrocellulose membranes (Supported Nitrocellulose Membrane 0.2 micron, Bio-Rad Laboratories Inc.) with Bio-Dot (Bio-Rad Laboratories Inc.) for 30 minutes. Next, these nanospheres were aspirated onto a nitrocellulose membrane, which was incubated with Blocking One for 1 hour at room temperature followed by washing in TBS-T three times. The membrane was incubated with anti-CXCR4 antibodies (200 ng/ml) at 4 ℃ overnight. After washing with TBS-T, the membrane was incubated with anti-mouse IgG HRP-conjugated antibodies in TBS-T (20 ng/ml) for 1 hour at room temperature. After washing with TBS-T, the membrane was incubated with ECL substrate and exposed using a Bio-Rad ChemiDoc XRS plus system.

### Evaluation of the binding affinity of MSC-GNS to SDF-1

SDF-1 (Human CXCL12/SDF-1β Protein (Fc Tag), Sino Biological Inc., Kanagawa, Japan) was labeled with biotin using the Biotin Labelling Kit-NH2 (Dojindo, Inc., Kumamoto, Japan) according to the manufacturer’s protocol. Twenty µg of MSC-GNS or GNS was incubated with the indicated concentrations of biotin-labeled SDF-1 at 37 °C for 1 hour. Next, the nanospheres were centrifuged at 14,000 ×g for 30 minutes three times. Then, they were added onto a nitrocellulose membrane using Bio-Dot, which was incubated with ELISA/ELISPOT Diluent (Thermo Fisher Scientific Inc.) for 1 hour, and washed with TBS-T three times. Next, the membrane was incubated with Pierce High Sensitivity NeutrAvidin-HRP (50 ng/ml; Thermo Fisher Scientific Inc.) in TBS-T for 1 hour. After washing with TBS-T, the membrane was incubated with ECL substrate and exposed using the Bio-Rad ChemiDoc XRS plus System.

### Collagenase-dependent degradation and controlled drug release assay

To evaluate the collagenase-dependent degradation of GNS, MSC-GNS or GNS at a final concentration of 200 μg/ml was incubated in PBS (+) solution with or without 0.05 U/ml collagenase D (Merck Co., Darmstadt, Germany) at 37 ℃. At different time points, the transmittance of the solutions at 500 nm was measured using a UV-visible spectrophotometer (nanoVette DU800, Beckman Coulter, Inc., Brea, CA, USA). The solution was lightly dispersed by tapping before measurement.

To evaluate the controlled drug release by a degradation-dependent mechanism, GNS-LSKL or MSC-GNS-LSKL was incubated in PBS (+) solution with or without 0.05 U/ml collagenase D in a 1.5 ml tube at 37 ℃. At different time points, the solution was centrifuged and then the supernatant was collected. The amount of LSKL in the supernatant was measured by HPLC. The calculation of LSKL release rates was performed according to the aforementioned method.

### Cellular uptake of nanospheres

RAW264.7 cells or MCT3T3-E1 cells were seeded onto 6 well polystyrene plates at a density of 1 × 10^5^ cells/well and cultured for 24 hours. Then, 20 μg/ml MSC-GNS_QD605_ or GNS_QD605_ was added and incubated for 6 hours with RAW264.7 cells or for 12 hours with MC3T3-E1 cells. Then, the cells were washed with PBS and collected by centrifugation at 200 ×g. The fluorescence signals in cells were analyzed by flow cytometry (BD FACSCount II, Becton Dickinson Inc., Franklin Lakes, NJ, USA) using Flow Jo software.

In addition, RAW264.7 cells or MCT3T3-E1 cells were seeded onto 24 well glass-bottom plates at a density of 2.5 × 10^4^ cells/well and cultured for 24 hours. Then, the cells were incubated with MSC-GNS_QD605_ or GNS_QD605_ under the same conditions as for the flow cytometric experiment. After washing with PBS, the nuclei were stained with Hoechst 33342 (Nacalai Tesque Inc.). Fluorescence signals were observed by confocal laser scanning microscopy (Olympus FV1000, Olympus Co., Tokyo, Japan).

### Mice and CCl_4_-induced liver fibrosis mouse model

Six-week-old male BALB/c mice were purchased from Japan SLC Inc. (Shizuoka, Japan). The animals were maintained on a standard food and water diet in a temperature-and light-controlled environment. All animal experiments were approved by the Kyoto University Animal Experimentation Committee and conducted in accordance with Regulations on Animal Experimentation at Kyoto University. To establish a mouse model of liver fibrosis, mice were intraperitoneally injected with olive oil with or without 20% carbon tetrachloride twice a week for a total of eight injections. Two days after the last injection, blood samples were collected and centrifuged at 800 ×g for 5 minutes to collect plasma. The levels of AST and ALT in the plasma were measured using a transaminase test (Fujifilm Wako Pure Chemical, Co., Osaka, Japan) to confirm the establishment of the mouse model of liver fibrosis. In addition, histological analysis was performed to confirm liver fibrosis. Briefly, mice were sacrificed, and their livers were fixed with 4% formaldehyde (MildformR 10N, Fujifilm Wako Pure Chemical Co.) overnight. The preparation of paraffin sections 4 µm thick and HE and MT staining were performed at the Kyoto Institute of Nutrition & Pathology, Inc. (Kyoto, Japan).

### Blood clearance of MSC-GNS after intravenous injection

One hundred µg of MSC-GNS_QD705_ or GNS_QD705_ was injected into the tail veins of normal mice. At the indicated time points, blood samples were collected from the tail vein, and plasma was collected by the centrifugation of blood at 800 ×g for 5 minutes. The fluorescence intensity of Qdots in plasma was measured with a spectrofluorometer. The obtained data were analyzed using R with the drc extension package.

### Tissue distribution of nanospheres after intravenous injection

Two days after the establishment of the mouse model of liver fibrosis, 100 µg of MSC-GNS_QD705_ or GNS_QD705_ was injected into the tail veins of the liver fibrosis model mice and control mice. Mice were sacrificed 3 or 24 hours after the injection, and their lungs, hearts, livers, spleens, kidneys, and blood from the inferior vena cava were collected. These organs were washed with saline, and fluorescent images were captured using an *in vivo* imaging system (IVIS Spectrum Imaging System, Caliper Life Sciences Inc., Hopkinton, MA, USA). The fluorescence intensity of the region of interest was analyzed by Living Image software (Caliper Life Sciences).

### Therapeutic effect of MSC-GNS on liver fibrosis

To establish the mouse model of liver fibrosis, mice were administered carbon tetrachloride intraperitoneally 12 times over 6 weeks. Mice were intravenously injected with MSC-GNS-LSKL (100 µg), GNS-LSKL (100 µg), an equal amount of LSKL, or saline (100 µl) six times over two weeks, starting 4 weeks after the initial carbon tetrachloride injection. Two days after the last treatment, mice were sacrificed, and their livers and blood from the inferior vena cava were collected. The plasma concentrations of AST and ALT were evaluated. Parts of extracted livers were evaluated histologically by HE staining and MT staining. For MT staining, the percentage of the blue area indicating fibrotic areas was quantified using ImageJ software.

## Statistical analysis

All quantitative data were expressed as the means ± standard deviation (SD). *t*-tests and one-way analysis of variance (ANOVA) followed by Tukey’s test were used for comparisons. Differences with a *p*-value <0.05 were considered statistically significant. (*: *p*<0.05, **: *p*<0.01, *#*: *p*<0.005, and *##*: *p*<0.0001). All statistical analyses were performed using GraphPad Prism 9.0 (GraphPad Software, Inc., La Jolla, CA, USA).

## Supplementary Materials

Materials and Methods Figs. S1 to S7

Tables S1

## Author Contribution

K.M. and M. A. designed and performed the all studies, and wrote the whole initial draft of this manuscript including supplementary information. K. S. and I. A. assisted with Cryo-EM observation. M.A. and Y. T. are the corresponding author supervised this project, and critically drafting this manuscript. All authors have read and approved the final manuscript.

## Funding

Grants-in-Aid for Scientific Research B (JP 23K25177 to M.A.) from the Japan Society for the Promotion of Science (JSPS).

MEXT Promotion of Distinctive Joint Research Center Program (JPMXP 0621467946).

## Acknowledgments

The authors thank Mr. Matsunaga, Instrumentation Center of The University of Kitakyushu, for the cryo-EM observation of nanoparticles using JEM-2100Plus. The authors also wish to thank Professor Y. Sasaki (Kyoto University) for help with electron microscopy and single particle analysis using an imaging flow cytometer. We thank J. Ludovic Croxford, PhD, from Edanz (https://jp.edanz.com/ac) for editing a draft of this manuscript.

## Abbreviations

ALT: alanine aminotransferase
AST: aspartate aminotransferase
α-SMA: α-smooth muscle actin
cGNS: cationized gelatin nanospheres
Cryo-EM: cryo-electron microscope
CXCR4: CXC-chemokine receptor 4
DDS: drug delivery system
GNS: gelatin nanospheres
HE staining: hematoxylin and eosin staining
HPLC: high performance liquid chromatography
LSKL: Leu-Ser-Lys-Leu-NH2
MSC: mesenchymal stem cells
MSC-GNS: mesenchymal stem cells membrane-coated gelatin nanospheres
MT staining: Masson’s trichrome staining
PLGA: poly lactic-co-glycolic acid
SDF-1: stromal cell-derived factor-1
TEM: transmission electron microscope
TGF-β: transforming growth factor-β
TSP-1: thrombospondin-1

