## Supplementary Information for "Mesenchymal Stem Cell Membrane Coating of Gelatin Nanospheres with Drug Release Ability for Inflammatory Tissue Targeting"

### Supplementary Materials
### for

### Mesenchymal Stem Cell Membrane Coating of Gelatin Nanospheres with Drug Release Ability for Inflammatory Tissue Targeting

Keisuke Moriyama[1], Mitsuru Ando[1*], Kazuo Sakurai[2], Isamu Akiba[3] Yasuhiko Tabata[4*]

[1] Laboratory of Biomaterials, Institute for Life and Medical Sciences, Kyoto University, 53 Kawara-cho Sohgoin, Sakyo-ku, Kyoto, 606-8507, Japan

[2] Department of Chemistry and Biochemistry, University of Kitakyushu, 1-1 Hibikino, Kitakyushu, Fukuoka, 808-0135, Japan

[3] Faculty of Environmental Engineering, the University of Kitakyushu, 1-1 Hibikino, Wakamatsu, Kitakyushu, Fukuoka 808-0135, Japan.

[4] Department of Plastic and Reconstructive Surgery, Graduate School of Medicine, Kyoto University, 54 Kawara-cho Sohgoin, Sakyo-ku, Kyoto, 606-8507, Japan

* : Corresponding author

### Materials and Methods

#### Preparation of cationized gelatin nanosphere (cGNS)

Gelatin (isoelectric point: pI 9.0, weight-averaged molecular weight: Mw 100,000) was kindly supplied from Nitta Gelatin Inc., Osaka, Japan. Cationized gelatin was prepared according to the slight modificated method previously reported.[S1] In briefly, 6.21 ml of ethylenediamine (Nacalai Tesque Inc., Kyoto, Japan) was added at 50-fold molar equivalent to the carboxyl group of gelatin into 50 ml of double-distilled water (DDW) containing 2.0 g of gelatin. The pH of solution was adjusted to 5.5 by adding 6 M hydrochloric acid (HCl) (Nacalai Tesque Inc., Kyoto, Japan.). 1-Ethyl-3-(3-(dimethylamino) propyl) carbodiimide hydrochloride (EDC) (Nacalai Tesque Inc., Kyoto, Japan) was added at 3-fold molar equivalent to the carboxyl groups of gelatin, followed by the addition of DDW into the solution to adjust the total volume of 100 ml. Then, the chemical introduction was allowed to proceed at 40 °C for 18 h under stirring, and then the resulting solution was dialyzed against DDW for 3 days at room temperature, followed by solution freeze-drying to obtain cationized gelatin. To determine the percentage of ethylenediamine introduced into the carboxyl groups of gelatin, the conventional 2,4,6-trinitrobenzenesulfonic acid (TNBS, FUJIFILM Wako Pure Chemical Corporation, Osaka, Japan) method was performed to be the percent introduced of 44.7%. Then, cGNS were prepared by a slight modification of the conventional coacervation method. [S1]

#### Western blot analysis of MSC-GNS

Samples were mixed with laemmli buffer containing 10% 2-mercaptoethanol and incubated at 70°C for 15 min. Then they were separated by SDS-PAGE under the reducing condition. Subsequently, bands were electrotransferred to a poly (vinylidene difluoride) PVDF membrane (Trans-Blot Turbo 0.2-micron PVDF Membrane, Bio-Rad Laboratories Inc.) at 4°C for 16 hours. A PVDF membrane was incubated with Blocking One (Nacalai Tesque Inc.) at room temperature for 1 hours and wash using Tris-buffered saline containing 0.02% Tween-20 (TBS-T) three times. Next, the membrane incubated with anti-CXCR4 antibodies (4G10, 200 ng/ml; Santa Cruz Biotechnology, Inc., Santa Cruz, CA, USA), anti-glyceraldehyde-3-phosphate dehydrogenase (GAPDH) antibodies (D16H11, 200 ng/ml; Cell Signaling Technology Inc., Danvers, MA, USA) or anti-CD47 antibodies (ab214453, 500 ng/ml; Abcam Ltd., Cambridge, UK) in TBS-T containing 10% Blocking One at 4°C overnight, respectively. After washing with TBS-T, the membrane was incubated with anti-mouse IgG HRP-conjugated antibodies (20 ng/ml; Cell Signaling Technology Inc., Danvers, MA, USA) for CXCR4 detection, anti-rabbit IgG HRP-conjugated antibodies (20 ng/ml; Cell Signaling Technology Inc.) for GAPDH detection, or anti-rabbit IgG HRP-conjugated antibodies for CD47 detection (20 ng/ml; Cell Signaling Technology Inc.) at room temperature for 1 hour, respectively. After washing with TBS-T, the membrane was incubated with the electrochemical luminescence (ECL) substrate (Pierce ECL Plus Western Blotting Substrate, Thermo Fisher Scientific Inc., Waltham, MA, USA) and exposed using Bio-Rad ChemiDoc XRS plus system (Bio-Rad Laboratories Inc.).

**Detection of LSKL peptide**

LSKL was evaluated by HPLC (Prominence LC-20AT; Shimazu Co., Ltd., Kyoto, Japan). A $C_{18}$ column was used for HPLC, and $H_2O$ containing 0.1% trifluoroacetic acid and acetonitrile (Nacalai Tesque Inc.) containing 0.1% trifluoroacetic acid (Nacalai Tesque Inc.) were used as mobile phases. The LSKL incorporating rate was calculated from the following equation.

$$LSKL\ incorporatin\ rate\ (\%) = \frac{The\ amount\ of\ LSKL\ added\ to\ GNS - The\ amount\ of\ LSKL\ in\ centrifugal\ supernatant}{The\ amount\ of\ LSKL\ added\ to\ GNS} \times 100$$

#### The measurement of cytotoxicity of MSC-GNS

Cell viability was evaluated by CCK-8 assay (Cell Count Reagent; Nacalai Tesque Inc.) according to the manufacturer's protocol. RAW 264.7 cells or MC3T3-E1 cells were seeded onto 96 well polystyrene plates at a density of 5,000 cells/100 μl RPMI 1640/well and were cultured for 24 hours. Then, the cells were incubated with MSC-GNS or GNS at the indicated concentrations for 24 hours. After incubation, 10 μl Cell Count Reagent was added to each well and incubated at 37°C for 3 hours. The absorbance at 450 nm was measured.

#### Establishment of mouse models of liver fibrosis

Six-week-old male Balb/c mice were purchased from Japan SLC Inc. (Shizuoka, Japan). The animals were maintained on a standard food and water diet in a temperature- and light-controlled environment. All animal experiments were approved by the Kyoto University Animal Experimentation Committee and conducted in accordance with Regulations on Animal Experimentation at Kyoto University. To establish mice model of liver fibrosis, mice were intraperitoneally injected with olive oil with or without 20% carbon tetrachloride twice a week for a total of 8 injections. Two days after the last injection, the blood samples were collected and centrifuged at 800×g for 5 minutes to collect the plasma. The levels of aspartate aminotransferase (AST) and alanine aminotransferase (ALT) in the plasma were measured by Transaminase test Wako (Fujifilm Wako Pure Chemical, Co., Osaka, Japan) to confirm the establishment of the mice model of liver fibrosis. In addition, the histological analysis was performed to confirm the liver fibrosis. Briefly, mice were sacrificed, and their livers were fixed with 4% formaldehyde (mildform R 10 N, Fujifilm Wako Pure Chemical Co.) overnight. The preparation of paraffin sections 4 μm thick and Hematoxylin and Eosin (HE) staining and Masson's trichrome (MT) staining were performed in Kyoto Institute of Nutrition & Pathology, Inc. (Kyoto, Japan).

#### Measurement of SDF-1 concentration

About 100 mg liver samples were collected and placed in 500 μl of cell lysis buffer (Nacalai Tesque Inc.) containing phosphatase and protease inhibitors. The liver samples were homogenized on ice for 5 minutes using Biomasher II (Nippi Co., Tokyo, Japan). The homogenized solution was centrifuged at 16,000×g for 15 minutes and the supernatants were collected. The concentration of SDF-1 in the supernatant and plasma were measured using the Mouse CXCL12/SDF-1 ELISA kit (Proteintech, Inc., Rosemont, IL, USA). The data are presented as mean ± s.d. ($n$ = 3)

**Table S1.** Hydrodynamic size and polydispersity index (PDI) of nanospheres mixed with MSC vesicles

| | Hydrodynamic size (nm) | PDI |
|---|---|---|
| MSC vesicles mixed with GNS | 288.3 ± 14.6 [a)] | 0.208 |
| MSC vesicles mixed with cGNS | 5574 ± 2077 | 0.671 |

a): mean ± SD

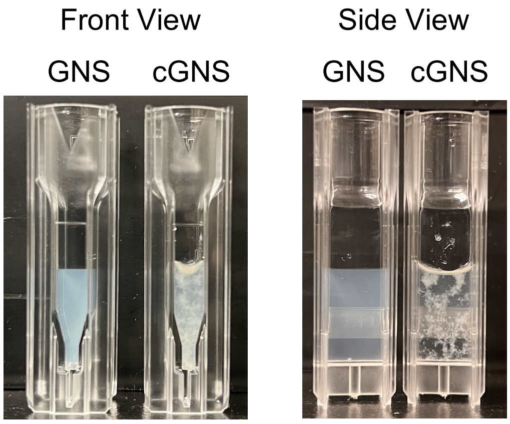


**Figure S1. Photograph of an aggregate formation after mixing cGNS with MSC vesicles.**

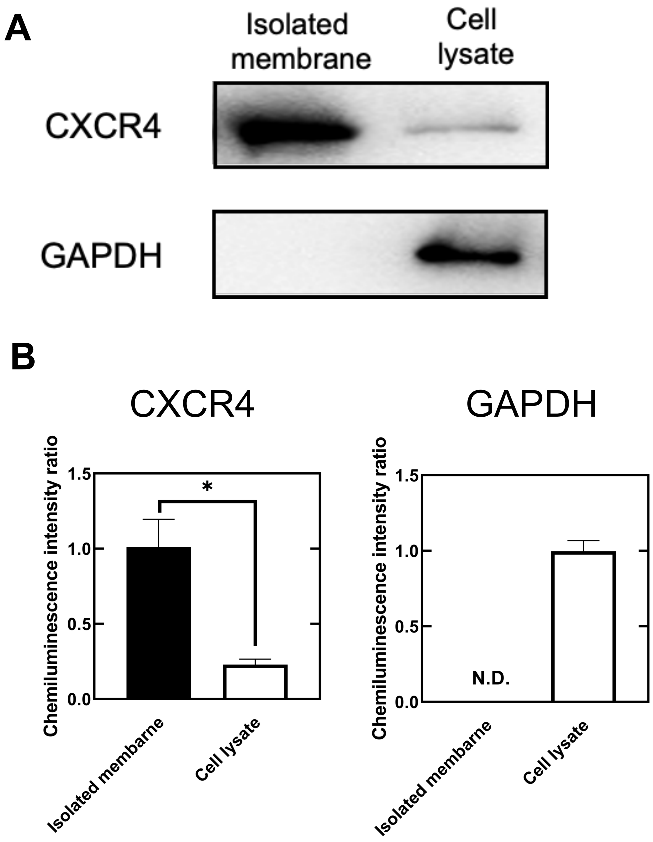


**Figure S2. Preparation and isolation of MSC membrane.** (A) Western blot analysis of CXCR4 and GAPDH in isolated membranes and cell lysates. (B) Semi-quantification of amount of CXCR4 and GAPDH, respectively. All results represent the average ± SD (n = 3). *p < 0.05, two-tailed Welch's t-test. N.D., not detected.

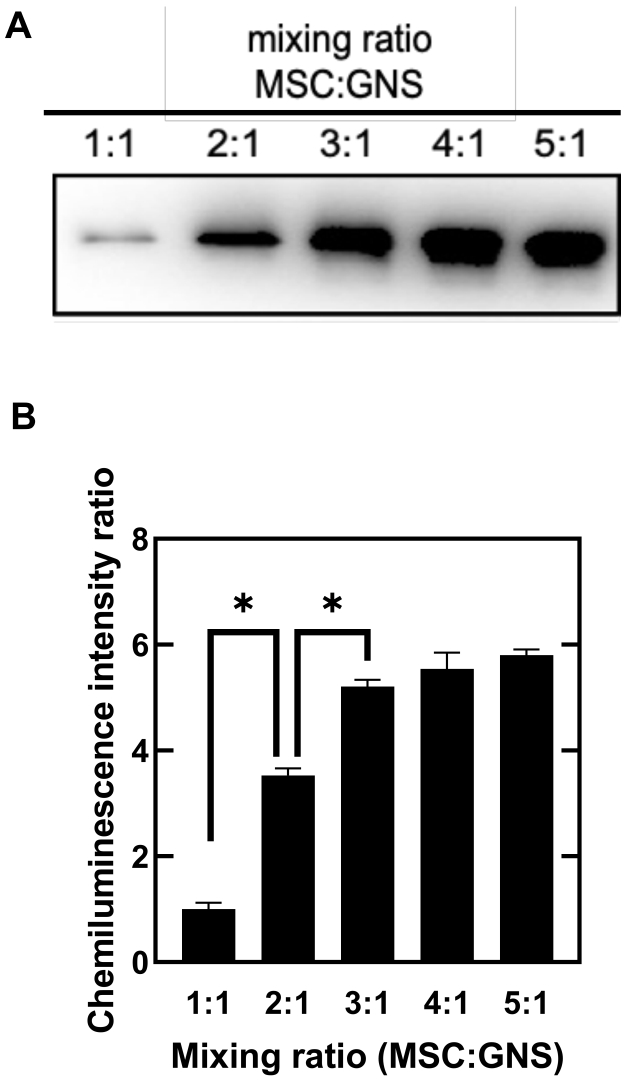


**Figure S3. Evaluation of the mixing ration between MSC membrane and GNS.** (A) Western blotting of CXCR4 in MSC-GNS at different mixing ratio. (B) Semi-quantification of amount of CXCR4 at different mixing ratio. All results represent the average ± SD (n = 3). One-way factorial ANOVA: *Adjusted $p < 0.05$, Tukey's post hoc test.

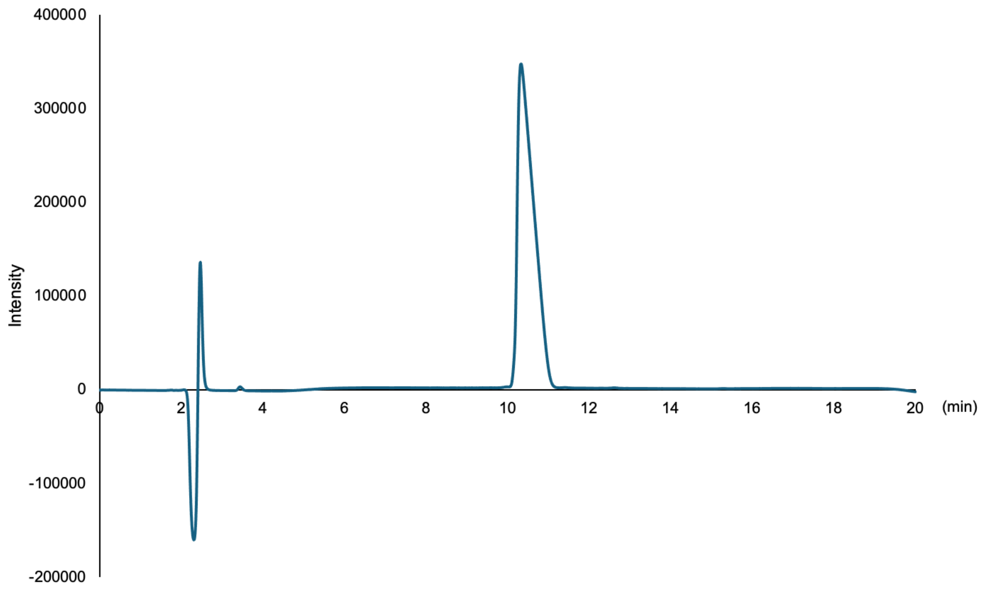


**Figure S4. Chromatograph of LSKL measured by HPLC.**

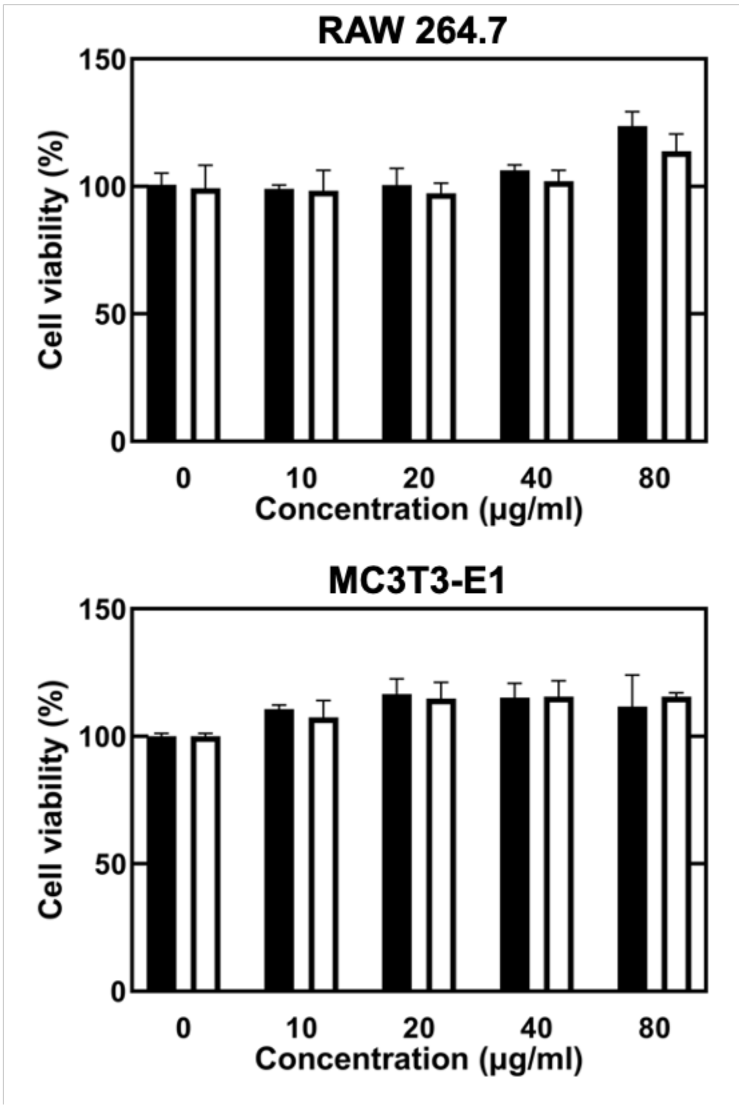


**Figure S5. Cytotoxicity of MSC-GNS.** Cell viability of RAW 264.6 (top) and MC3T3-E1 cells (bottom) 24 hr after incubation with MSC-GNS or GNS at different concentration. MSC-GNS (■) and GNS (□). All of the data are presented as mean ± s.d. (n = 3)

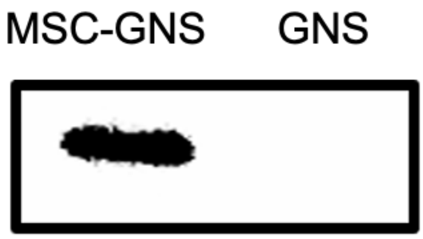


**Figure S6. Presence of CD47 on MSC-GNS and GNS by western blotting.**

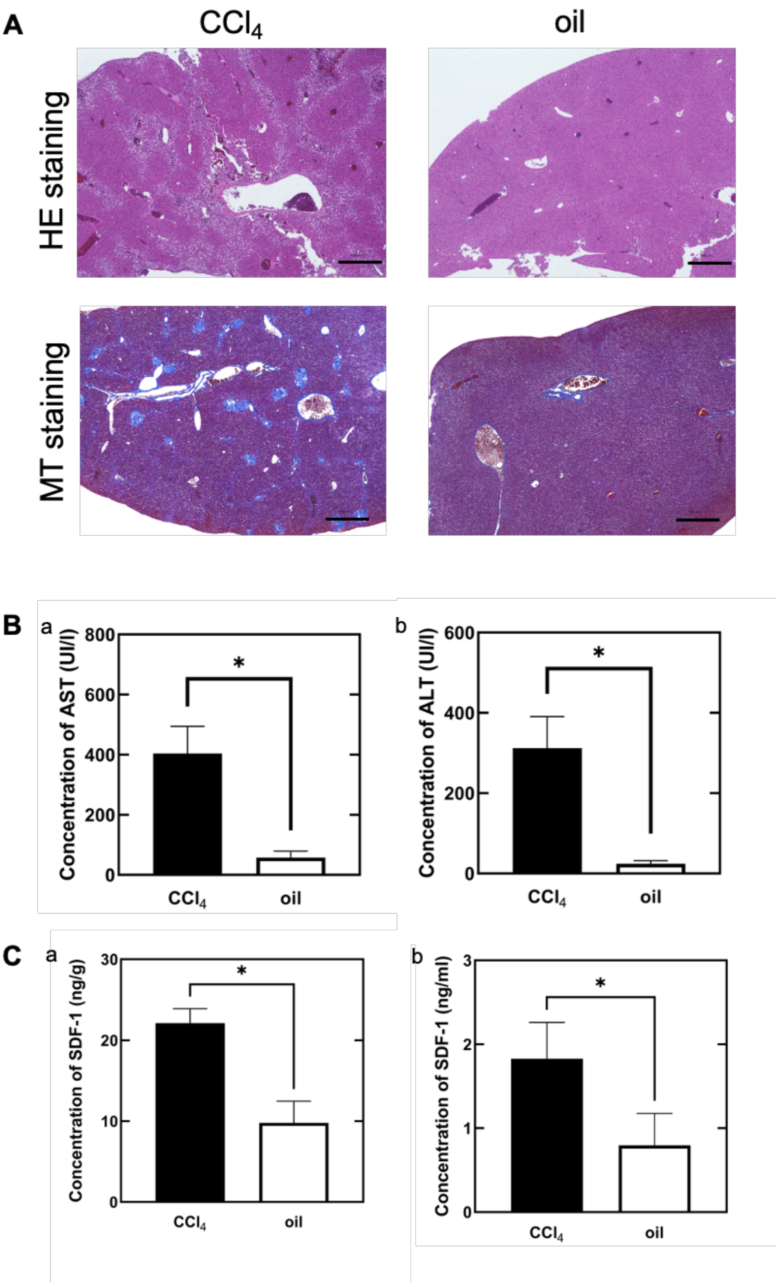


**Figure S7. Establishment of mouse models of liver fibrosis.** (A) Microscopic images of Hematoxylin and Eosin (HE) (top) and Masson trichrome (MT) staining (bottom) of liver of mice after intraperitoneal injection of carbon tetrachloride and olive oil. The scale bar is 500 μm. (B) AST (a) and ALT (b) levels in plasma of mice after intraperitoneal injection of carbon tetrachloride and olive oil, respectively. All results represent the average ± SD ($n = 3$). *$p < 0.05$, two-tailed Welch's t-test. N.D., not detected. (C) Expression levels of SDF-1 in liver (a) and plasma (b) of mice after intraperitoneal injection of carbon tetrachloride and olive oil. All of the data are presented as mean ± s.d. ($n = 3$) *$p < 0.05$, two-tailed Welch's t-test.
